# Sex and genetic background influence intravenous oxycodone self-administration in the Hybrid Rat Diversity Panel

**DOI:** 10.1101/2024.10.03.615948

**Authors:** Eamonn P. Duffy, Jack O. Ward, Luanne H. Hale, Kyle T. Brown, Andrew J. Kwilasz, Erika A. Mehrhoff, Laura M. Saba, Marissa A. Ehringer, Ryan K. Bachtell

## Abstract

Opioid Use Disorder (OUD) is an ongoing worldwide public health concern. Genetic factors contribute to multiple OUD-related phenotypes, such as opioid-induced analgesia, initiation of opioid use, and opioid dependence. Here, we present findings from a behavioral phenotyping protocol using male and female rats from 15 genetically diverse inbred strains from the Hybrid Rat Diversity Panel (HRDP). We used a self-administration paradigm to measure the acquisition of oxycodone intake during ten 2-hour sessions and escalation of oxycodone use during ten 12-hour sessions. During both the acquisition and escalation phases of self-administration, we observed that genetic background and sex influence oxycodone intake. The heritability of oxycodone intake phenotypes ranged between 0.26 to 0.54, indicating that genetic background plays a major role in the variability of oxycodone consumption. Genetic background and sex also influenced additional phenotypes recorded during oxycodone self-administration including lever discrimination and timeout responding. The genetic contribution to these traits was slightly more moderate, with heritability estimates ranging between 0.25 to 0.42. Measures of oxycodone intake were highly positively correlated between acquisition and escalation phases. Interestingly, the efficacy of oxycodone analgesia was positively correlated with oxycodone intake during the escalation phase, indicating that the initial behavioral responses to oxycodone may predict self-administration phenotypes. Together, these data demonstrate that sex and genetic background are major contributors to oxycodone self-administration phenotypes.

## 1. Introduction

Approximately 6.7-7.6 million individuals 12 years or older are estimated to have opioid use disorder (OUD) in the United States, and the opioid epidemic continues to pose a major public health issue (1). Prescription opioids have long been considered the gold standard for pain management (2,3). However, long-term use of these drugs confers an increased risk of several adverse health outcomes including tolerance, OUD, and overdose (4–6). Not all individuals who use prescription opioids develop signs of misuse, physical dependence, or OUD suggesting that some individuals may be more vulnerable to problematic use (7,8). Examining how genetic variability contributes to differences in the initiation of recreational opioid use and the progression to compulsive opioid use will be essential for understanding the biological underpinnings of OUD.

Heritability estimates for OUD-related traits range between 0.31 and 0.54 suggesting that genetic factors contribute to the risk of developing OUD in humans (9–11). Genome-wide association studies (GWAS) have had limited success in identifying specific genetic variants that contribute to the liability for developing OUD (12). Several GWAS have implicated a single nucleotide polymorphism (SNP) in *OPRM1* that confers a heightened risk of developing OUD, but few novel SNPs have been identified (13). The cross-sectional nature of most GWAS makes it difficult to study the genetic factors contributing to specific stages of OUD. Recent studies have tried to address this limitation by studying opioid misuse or problematic opioid use (POU), which may be analogous to the initial stages of OUD (14). POU is genetically correlated with OUD and opioid dependence, but there is little to no overlap in the genetic variants that confer heightened risk for POU or OUD (14). Although POU may contribute to the development of OUD, the progression between these phenotypes remains unclear. Rodent models can more precisely measure the initiation of opioid use and the shift to the compulsive-like escalation of opioid intake while also controlling for genetic and environmental factors.

Gender and sex also play a role in prescription opioid use and the development of OUD (15–19). Men display higher rates of nonmedical prescription opioid use and are at higher risk of prescription opioid overdose mortality compared to women (16,20). Human genetics studies have implicated candidate genes that may increase susceptibility to opioid dependence in a sex-specific manner (21,22). Unfortunately, sex differences or gene-by-sex interactions are not commonly studied or reported in GWAS of opioid-related traits (23–29). Many questions remain about how interactions between genetic background and sex affect opioid consumption and other related traits.

Rodent models can be effectively used to study how interactions between genetic background and sex influence opioid-related behaviors. The Hybrid Rat Diversity Panel (HRDP) is comprised of nearly 100 genetically divergent inbred rat strains, including two recombinant inbred panels and several divergent classic inbred strains (30). Due to the high genetic diversity between strains, this model is especially well-suited for examining genetic contributions to complex behavioral traits. Previous work from our lab demonstrated that both sex and genetic background influence behavioral measures of oxycodone-induced analgesia in HRDP strains (31). Oral and intravenous (i.v.) oxycodone self-administration has been reported in select strains from the HRDP, and the occurrence of sex differences in oxycodone intake is dependent on genetic background (32,33). Here, we used an i.v. self-administration paradigm to examine how interactions between genetic background and sex shape the acquisition and escalation of oxycodone self-administration phenotypes across 15 HRDP strains.

## 2. Methods

### 2.1 Subjects

Adult (PND 60+) male and female rats (n=266) from 15 HRDP strains were used for these experiments. Two strains (F344/NCrl and WKY/NCrl) were obtained from Charles River Laboratories, and the remaining 13 strains were obtained from the Medical College of Wisconsin (provided by Dr. Melinda Dwinell, R24OD024617). Strains and sample sizes are described in **Table 1**, and the attrition rate is depicted in **Supplemental Figure 1**. Upon arrival, rats were allowed to habituate for at least one week. Before intravenous surgery and self-administration procedures, all animals were tested for somatosensory and analgesia-related measures, as described in the **Supplemental Methods** and previously (31). Rats were single-housed in a temperature (22 °C) and humidity (40%) controlled animal vivarium on a 12-h light/12-h dark cycle. Rats were provided with standard rat chow (Teklad 2918, Envigo, Indianapolis, IN) and water *ad libitum*. Animals were tested in 22 cohorts (n=15-24/cohort). The ACI/EurMcwi strain was used as a control strain in all batches obtained from the Medical College of Wisconsin to allow for the detection and minimization of batch effects. This strain was chosen due to preliminary studies identifying that the ACI/EurMcwi strain displays robust behavioral responses in all outcome measures. All procedures were conducted in accordance with institutional guidelines outlined by the Institutional Animal Care and Use Committee at the University of Colorado Boulder, an Association for Assessment and Accreditation of Laboratory Animal Care International accredited institution.

**Table 1:**
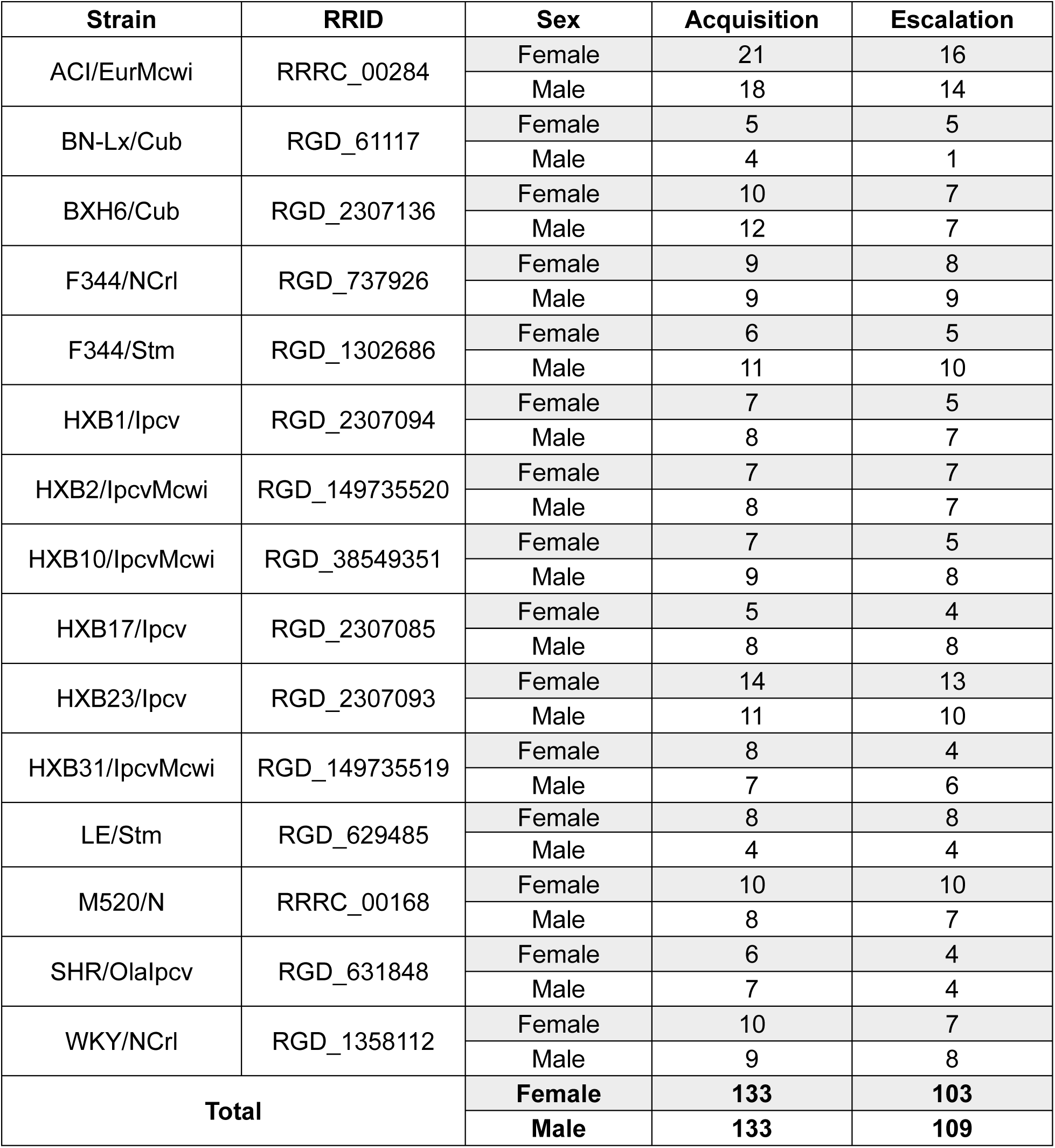
Sample size per strain and sex during the acquisition and escalation phases. Due to the low sample size in BN-Lx/Cub males, this strain was eliminated from statistical analyses during the escalation phase.

| Strain | RRID | Sex | Acquisition | Escalation |
| --- | --- | --- | --- | --- |
| ACI/EurMcwi | RRRC_00284 | Female | 21 | 16 |
|  |  | Male | 18 | 14 |
| BN-Lx/Cub | RGD_61117 | Female | 5 | 5 |
|  |  | Male | 4 | 1 |
| BXH6/Cub | RGD_2307136 | Female | 10 | 7 |
|  |  | Male | 12 | 7 |
| F344/NCrl | RGD_737926 | Female | 9 | 8 |
|  |  | Male | 9 | 9 |
| F344/Stm | RGD_1302686 | Female | 6 | 5 |
|  |  | Male | 11 | 10 |
| HXB1/lpcv | RGD_2307094 | Female | 7 | 5 |
|  |  | Male | 8 | 7 |
| HXB2/lpcvMcwi | RGD_149735520 | Female | 7 | 7 |
|  |  | Male | 8 | 7 |
| HXB10/lpcvMcwi | RGD_38549351 | Female | 7 | 5 |
|  |  | Male | 9 | 8 |
| HXB17/lpcv | RGD_2307085 | Female | 5 | 4 |
|  |  | Male | 8 | 8 |
| HXB23/lpcv | RGD_2307093 | Female | 14 | 13 |
|  |  | Male | 11 | 10 |
| HXB31/lpcvMcwi | RGD_149735519 | Female | 8 | 4 |
|  |  | Male | 7 | 6 |
| LE/Stm | RGD_629485 | Female | 8 | 8 |
|  |  | Male | 4 | 4 |
| M520/N | RRRC_00168 | Female | 10 | 10 |
|  |  | Male | 8 | 7 |
| SHR/OlaIpcv | RGD_631848 | Female | 6 | 4 |
|  |  | Male | 7 | 4 |
| WKY/NCrl | RGD_1358112 | Female | 10 | 7 |
|  |  | Male | 9 | 8 |
| Total |  | Female | 133 | 103 |
|  |  | Male | 133 | 109 |

### 2.2 Surgery

Chronic indwelling intra-jugular catheters were implanted using a modified procedure described previously (O’Neill et al., 2012). Animals were anesthetized (2-4% isoflurane), and intravenous catheters were implanted in the right jugular vein under aseptic conditions. Once the vein was isolated, it was punctured with a 22-gauge needle and the tubing was inserted and secured with suture thread. Sterile catheters (Access Technologies, Skokie, IL) consisted of 14 cm of polyurethane tubing (0.51 mm inner diameter, 1.12 mm outer diameter) secured to a 22-gauge back mount pedestal (Protech International). Immediately prior to surgery, animals were administered the analgesic carprofen (5 mg/kg, s.c.) and antibiotic enrofloxacin (5 mg/kg, s.c.). Animals received daily subcutaneous injections of carprofen (5 mg/kg) for two days following surgery. Catheters were flushed daily with 0.2 mL heparinized (20 IU/ml) bacteriostatic saline containing gentamicin sulfate (0.33 mg/ml; Hospira). Catheter patency was evaluated using 0.2 mL propofol (10 mg/mL propofol, Sagent) solution before the acquisition phase of oxycodone self-administration, before the escalation phase, and after the final session of the escalation phase.

### 2.3 Oxycodone self-administration

Self-administration procedures were conducted in operant chambers (Med Associates, St. Albans, VT) containing two retractable levers (a drug-paired left lever and an inactive right lever), stimulus cue lights above each lever, and a sound-attenuating fan. The acquisition and escalation of oxycodone intake was measured using a two-phase self-administration paradigm (**Figure 1A**). All self-administration sessions began at the start of the dark phase of the light/dark cycle. During the acquisition phase, rats were trained to self-administer oxycodone (Oxycodone HCl; B&B pharmaceutical, Englewood, CO) on a fixed-ratio 1 (FR1) reinforcement schedule in ten 2-hour “short-access” (ShA) sessions. Responses on the drug-paired lever resulted in an oxycodone infusion (0.15 mg/kg/infusion) delivered over 5-s and a cue light (7.5-W white light, 20-s). Each oxycodone infusion was followed by a 20-s timeout period (TO20). Responses on the inactive lever were recorded but had no consequences. After the acquisition phase, rats were tested under a progressive ratio (PR) schedule to evaluate motivation to obtain oxycodone. Due to a procedural error during the PR sessions, the data are unusable and not shown. Rats were then transitioned to the escalation phase that included ten 12-hour “long-access” (LgA) sessions under the same FR1:TO20 schedule of reinforcement (35).

**Figure 1.**
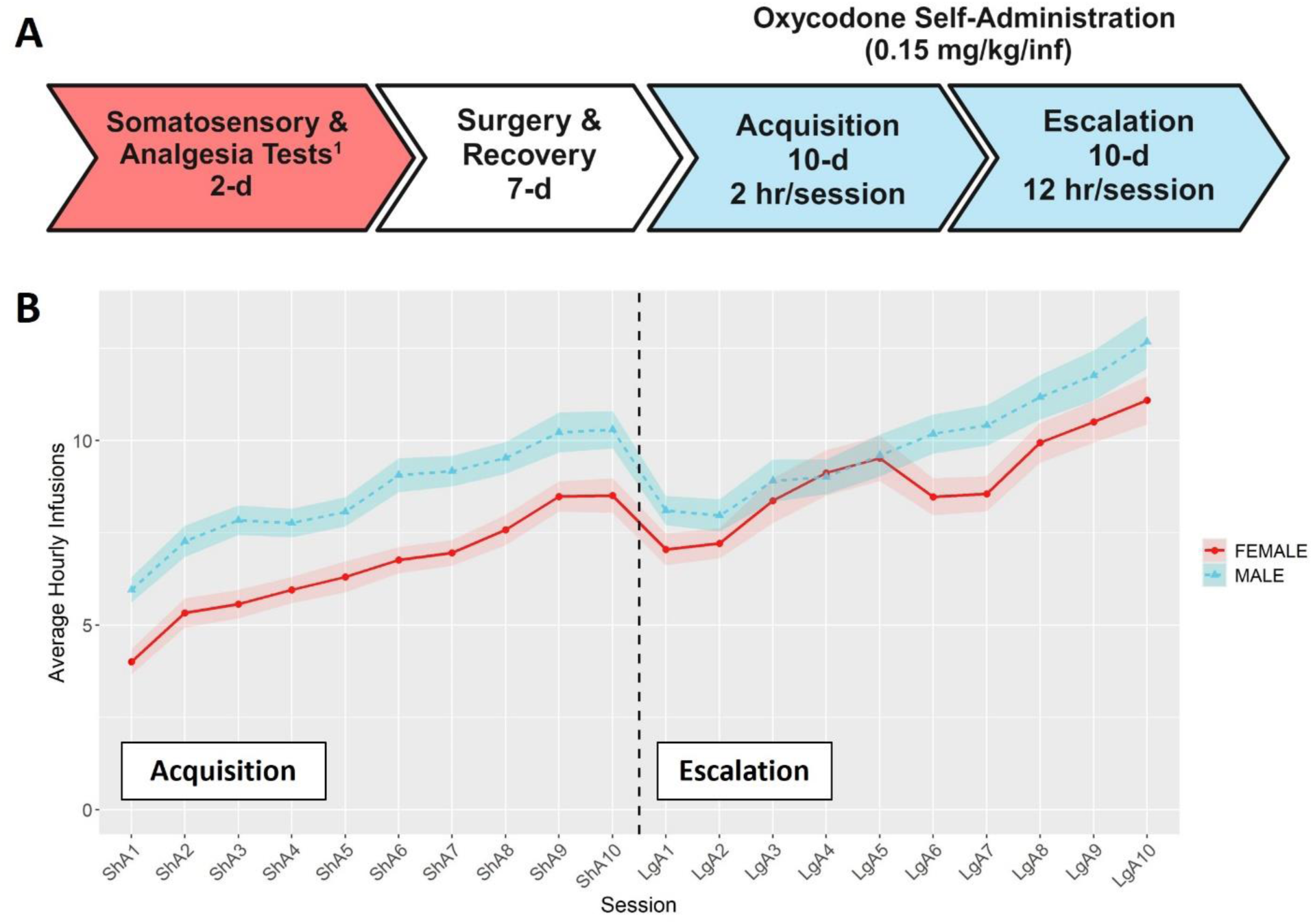
Overview of self-administration procedures. **(A)** Timeline for the behavioral phenotyping protocol. ^1^Results for somatosensory and analgesia tests are presented in (31). Timeline was create using BioRender.com **(B)** Hourly oxycodone intake (mean ± SEM) in male and female rats throughout the acquisition and escalation phases. The vertical dashed line represents the transition between phases. Both sexes increased their hourly oxycodone intake throughout both phases, and hourly intake was influenced by both sex and session.

### 2.4 Behavioral Phenotypes

#### Average Hourly Intake

The average hourly intake was calculated by dividing the number of infusions within a session by the duration of that session (hr). This measure was used to normalize the oxycodone intake across the acquisition (ShA; 2-hr) and escalation (LgA; 12-hr) sessions and depict the overall pattern of self-administration across both phases of self-administration.

#### Oxycodone Intake

Oxycodone intake was evaluated across different levels of detail. The most detailed and granular measure includes the number of oxycodone infusions per session. This was used to depict session-level data and describe the overall pattern of self-administration. Oxycodone intake was also evaluated by averaging oxycodone infusions across “early” (first three sessions) and “late” (last three sessions) time points. This measure was used to describe how oxycodone intake changes during the acquisition and escalation phases. Oxycodone intake at the early and late time points was also used to calculate a difference score for each rat as follows:

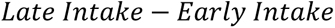

The broadest measure of oxycodone intake was the total oxycodone dose. This was calculated by adding the cumulative oxycodone dose received throughout acquisition or escalation.

#### Lever Discrimination

This phenotype was used to evaluate the ability of the rats to learn to discriminate between the drug-paired and inactive levers across the acquisition and escalation phases. We calculated a lever discrimination score during the early and late time points for each phase as follows:

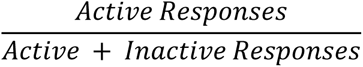

#### Normalized Timeout Responding

Timeout responding was evaluated as a measure of perseverative responding during periods when oxycodone is unavailable. Because animals that self-administer more oxycodone have more opportunities for timeout responding, the number of timeout responses was normalized to the number of drug infusions using the following equation:

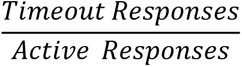

### 2.5 Broad-Sense Heritability Estimates

The coefficient of determination (R^2^) from a 2-way ANOVA with strain and sex as the predictors was used to estimate the broad-sense heritability (H^2^) for the following behavioral phenotypes during the acquisition and escalation phases: total oxycodone intake, early and late oxycodone intake, change in oxycodone intake, early and late lever discrimination, early and late normalized timeout responses.

### 2.6 Statistical Analysis

All data are presented as mean ± SEM. Longitudinal data for hourly intake or intake per session was analyzed using a repeated measures ANOVA with session as the within-subject factor and strain and sex as the between-subject factors. Oxycodone intake, lever discrimination, and normalized timeout responding were analyzed with repeated measures ANOVAs with timepoint (early or late) as the within-subject factor and strain and sex as the between-subject factors. Results from repeated measures ANOVAs are reported in **Supplemental Table 1.** Two-way ANOVAs were run to test the effects of strain, sex, and their interaction for total oxycodone intake and the change in oxycodone intake. Results from two-way ANOVAs are presented in **Supplemental Table 2**. To examine sex differences in total oxycodone intake during acquisition and escalation, a simple main effects analysis was performed in which Student’s t-tests were used to test the significance of sex within each strain. Pearson correlations were used to evaluate the relationship between oxycodone analgesia and self-administration phenotypes, using strain means calculated within each sex for each behavioral phenotype. Pairwise correlations between oxycodone behavioral phenotypes are presented in **Supplemental Table 3.**

Outliers were identified and winsorized prior to statistical analysis, as described in the **Supplemental Methods**. Missing data (< 2% of self-administration session data) was imputed as described in the **Supplemental Methods.** Statistical significance was set as p < 0.05. For the sex differences analysis, a Bonferroni correction was applied based on the number of tests for 15 strains during acquisition (p < 0.003) and 14 strains during escalation (p < 0.004). All data analysis was conducted in R v4.3.2.

## 3. Results

### 3.1 Oxycodone self-administration in males and females

Male and female rats from 15 HRDP strains underwent a behavioral protocol designed to measure the acquisition and escalation of i.v. oxycodone self-administration (**Figure 1A**). The average hourly oxycodone intake was calculated to describe and compare the general pattern of oxycodone consumption in males and females during acquisition and escalation sessions collapsed across strains. We observed a general pattern of increasing oxycodone intake across the entire procedure **(Figure 1B**, F_6.37,1368.52_ = 45.55, p < 0.001). Both males and females increased their oxycodone intake throughout the procedure, with males generally showing higher intake across sessions (F_1,215_ = 9.82, p = 0.002). Next, we analyzed how the interaction between genetic background and sex influenced various aspects of self-administration.

### 3.2 Oxycodone intake

To assess strain and sex differences in the acquisition of oxycodone self-administration rats were trained to self-administer oxycodone in 10 daily short-access sessions. The general pattern of oxycodone intake per session during the acquisition phase **(Figure 2)** was shaped by a strain-sex interaction (F_14,236_ = 3.24, p < 0.001) and a strain-session interaction (F_76.12,1283.21_ = 3.11, p < 0.001). Males and females from a few strains (e.g., HXB2/IpcvMcwi) displayed divergent patterns of oxycodone consumption, with males having greater intake than females during acquisition. Strains also differed in the general pattern of self-administration over time. Some strains (e.g., HXB23/Ipcv and HXB31/IpcvMcwi) displayed relatively stable responding while others (e.g., HXB1/Ipcv) rapidly increased oxycodone intake across the acquisition phase. To simplify the strain and sex comparisons, we calculated the total oxycodone dose consumed during the acquisition phase. **Figure 3** illustrates that total oxycodone intake was shaped by a strain-sex interaction (F_14,236_ = 3.24, p < 0.001). Males from the HXB2/IpcvMcwi strain consumed more oxycodone than females (t_8.29_ = −6.59, p < 0.001). Males from the ACI/EurMcwi (t_29.71_ = −2.89, p = 0.007) and WKY/NCrl (t_15.73_ = −2.56, p = 0.02) strains also displayed increased oxycodone intake compared to their female counterparts, but these sex differences did not survive multiple testing correction.

**Figure 2.**
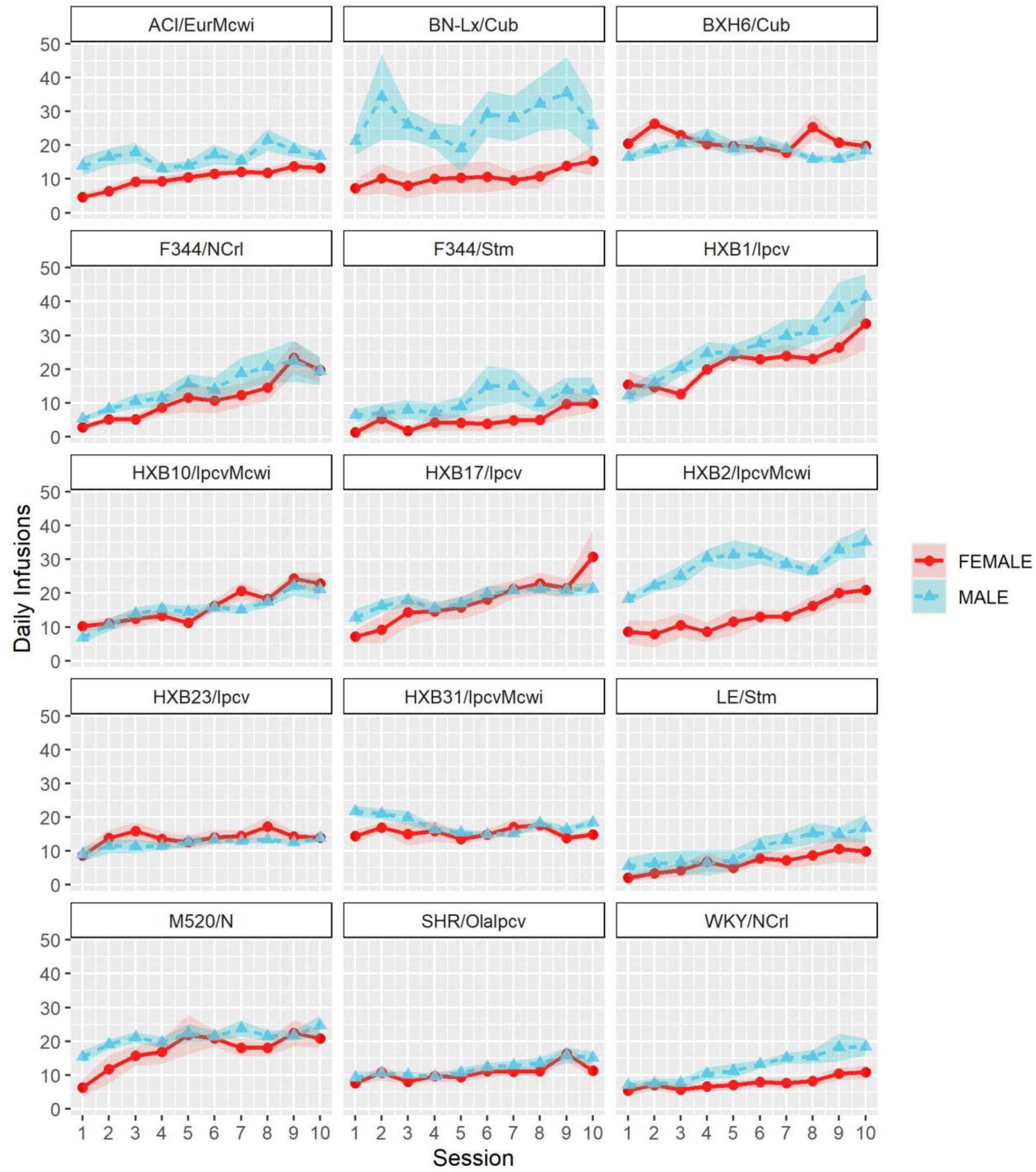
Acquisition of oxycodone intake is influenced by strain and sex. Daily oxycodone intake (mean ± SEM) was measured during 2-hr sessions and is plotted in separate graphs for each strain. Patterns of oxycodone intake were shaped by a strain-sex interaction and a strain-session interaction.

**Figure 3.**
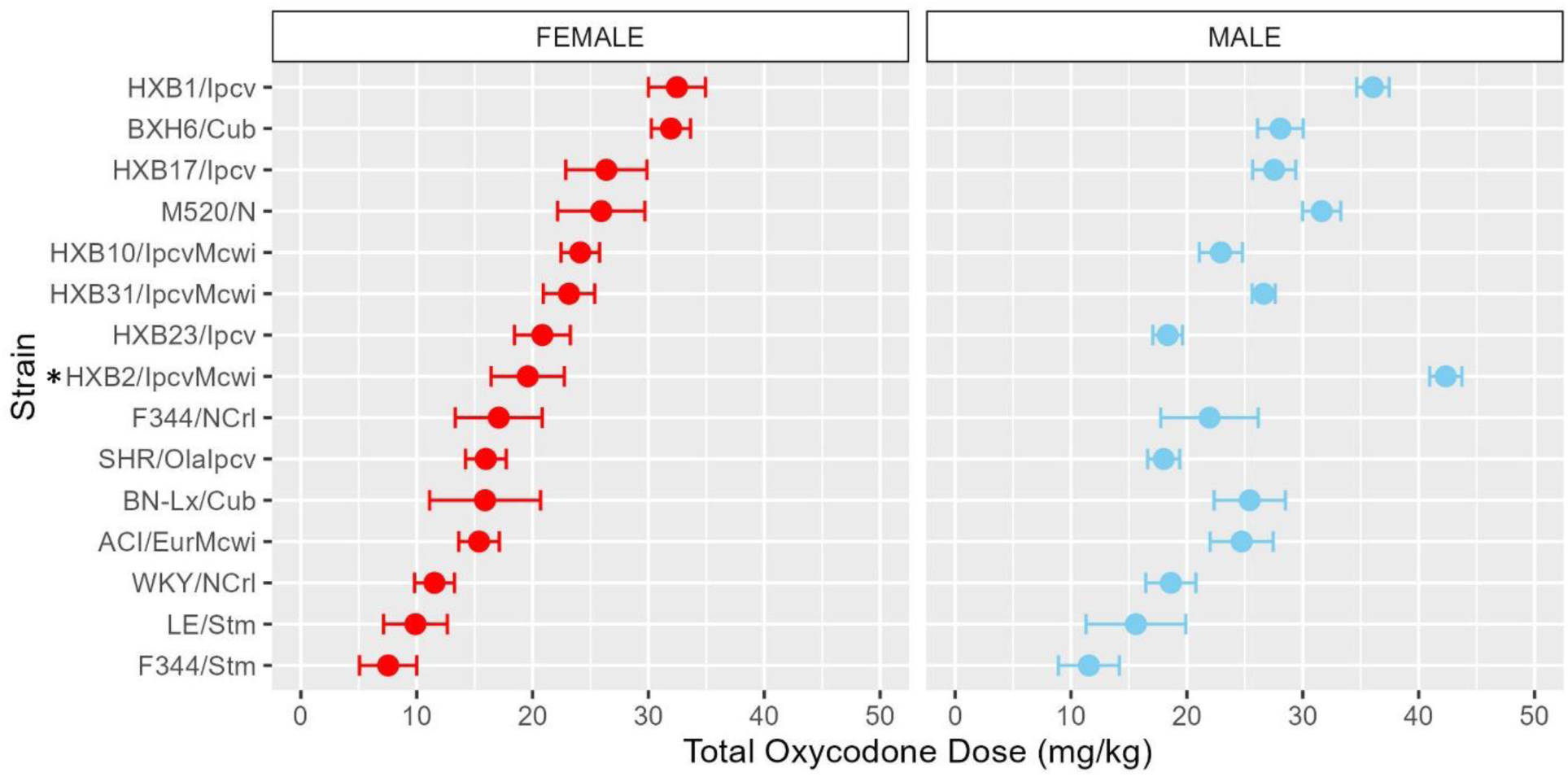
Total oxycodone intake during acquisition is influenced by strain and sex. Daily oxycodone infusions were summed to calculate the total oxycodone dose consumed during the acquisition phase. The mean ± SEM for each strain is plotted for females (left) and males (right). Strains for both females and males are displayed in order of increasing total oxycodone dose in females. The interaction between genetic background and sex influence the total amount of oxycodone self-administered during acquisition. * denotes sex differences within strain.

To further characterize how patterns of oxycodone consumption change across the acquisition period the mean intake during the first three acquisition sessions (“early”) and last three acquisition sessions (“late”) was calculated **(Supplemental Figure 2)**. Within these windows, oxycodone intake was shaped by a strain-sex interaction (F_14,236_ = 3.66, p < 0.001) and a strain-timepoint interaction (F_14,236_ = 6.99, p < 0.001). To further explore these strain differences, the change in intake was calculated as the difference in intake between early and late acquisition **(Figure 4)**. Genetic background influenced the change in oxycodone intake across acquisition, as indicated by a main effect of strain (F_14,236_ = 6.97, p < 0.001). Although most strains increased oxycodone intake, some strains (e.g., BXH6/Cub, HXB31/IpcvMcwi) showed a minimal change in intake during the acquisition phase.

**Figure 4.**
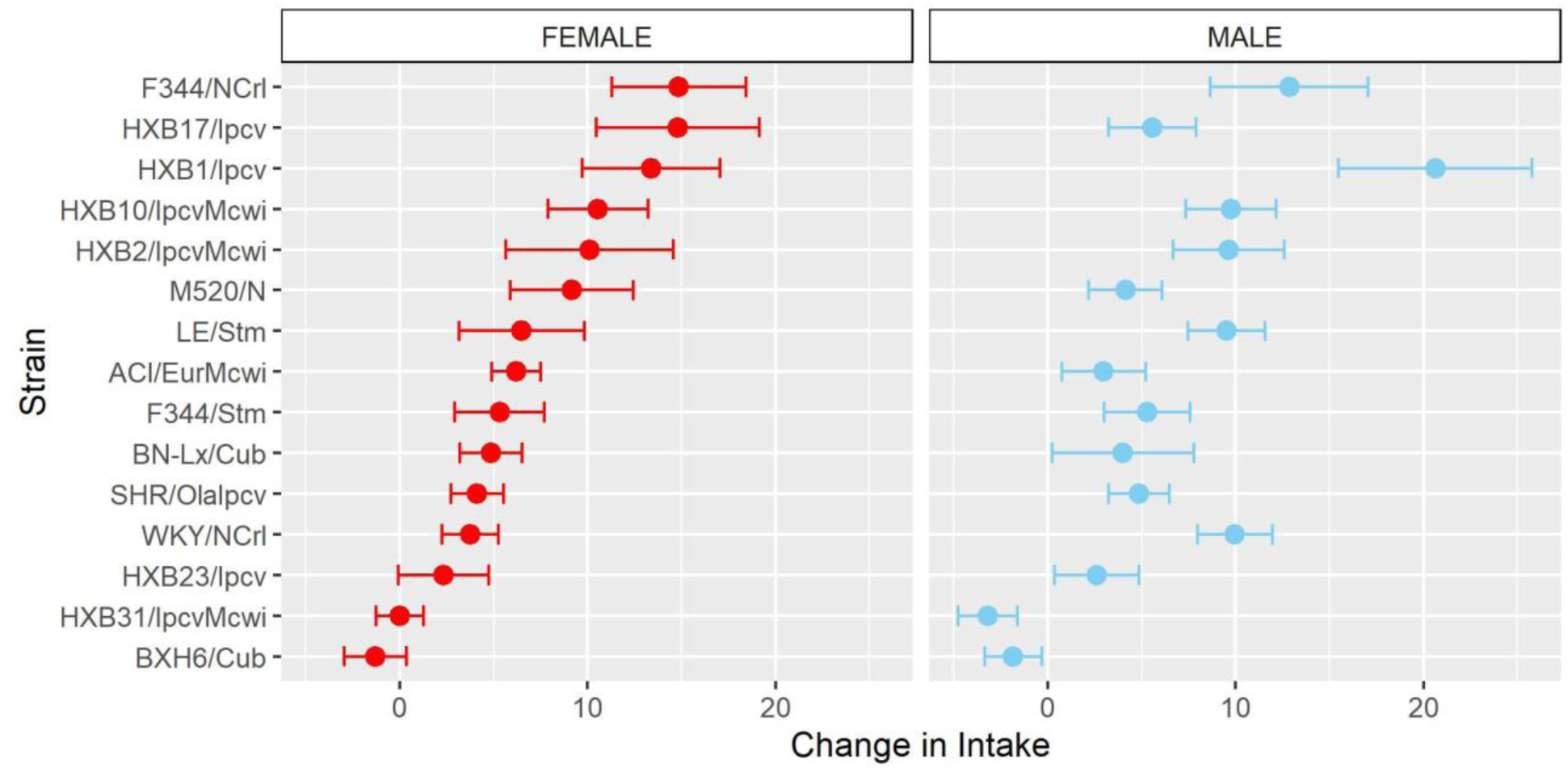
Change in oxycodone intake during acquisition is influenced by strain. The change in oxycodone intake was determined by calculating the difference between early and late acquisition sessions. The mean ± SEM for each strain is plotted for females (left) and males (right). Strains for both females and males are displayed in order of increasing change in females. Strains differed in their change in oxycodone intake.

Following acquisition, rats underwent 10 daily long-access sessions to measure the escalation of oxycodone self-administration. The BN-Lx/Cub strain is included in **Figure 5** to qualitatively describe their pattern of self-administration but was excluded from statistical analysis due to the low sample size in the males (n=1). Throughout escalation, oxycodone intake **(Figure 5)** was influenced by a strain-sex (F_13,184_ = 4.44, p < 0.001) and a strain-session (F_70.49,997.76_ = 2.33, p < 0.001) interaction, suggesting that there is a complex interplay between how genetic background and sex influence the escalation of oxycodone intake. To simplify this interaction, the total oxycodone dose during the escalation phase was calculated **(Figure 6)**. Here, we again see that sex and strain contribute to overall oxycodone intake during the escalation sessions as indicated by a strain-sex interaction (F_13,184_ = 4.44, p < 0.001). A striking difference was observed in HXB2/IpcvMcwi males that self-administered significantly more oxycodone than females (t_11.33_ = −5.74, p = 0.001). Nominal sex differences were observed in the WKY/NCrl strain (t_7.32_ = −2.73, p = 0.03) but did not survive multiple testing correction.

**Figure 5.**
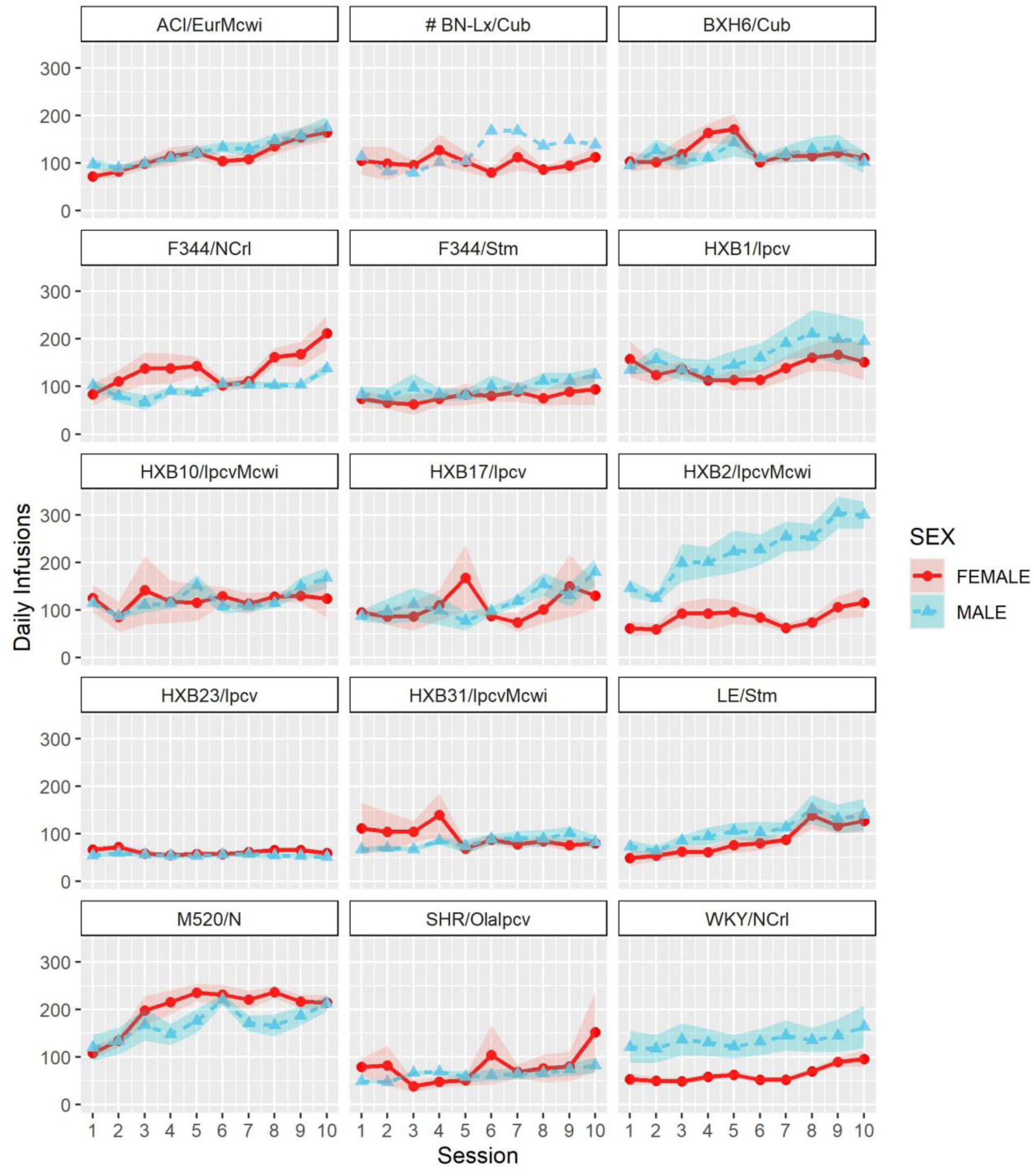
Escalation of oxycodone intake is influenced by strain and sex. Daily oxycodone intake (mean ± SEM) was measured during 12-hr sessions and is plotted in separate graphs for each strain. Oxycodone intake was influenced by strain-sex and strain-session interactions. ^#^ The BN-Lx/Cub strain is included for qualitative comparison but was excluded from statistical analyses due to low sample size resulting from attrition.

**Figure 6.**
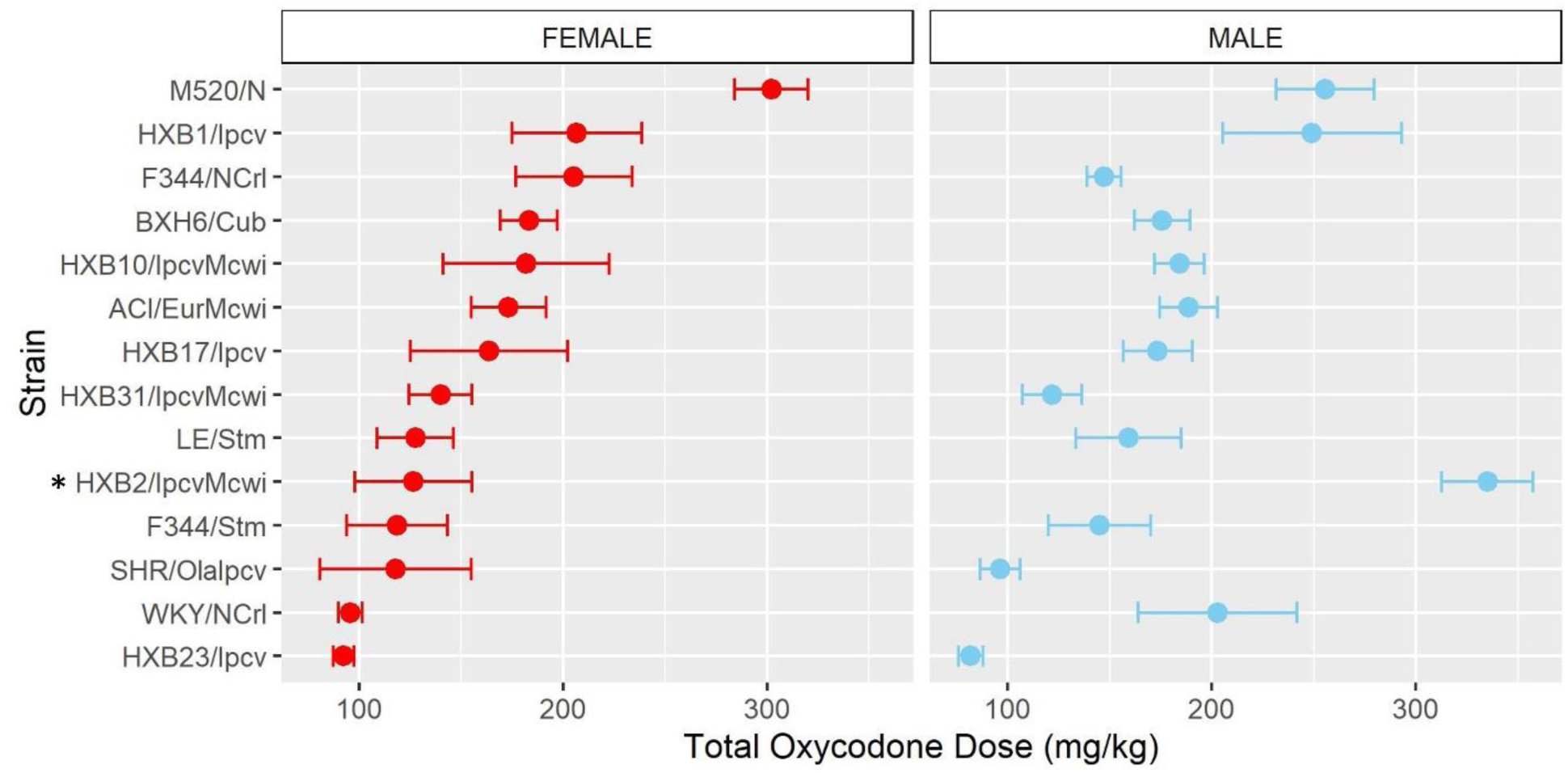
Total oxycodone intake during escalation was influenced by strain and sex. Daily oxycodone infusions were summed to calculate the total oxycodone dose consumed during the escalation phase. The mean ± SEM for each strain is plotted for females (left) and males (right). Strains for both females and males are displayed in order of increasing total oxycodone dose in females. The total amount of oxycodone self-administered during escalation was shaped by a strain-sex interaction. * denotes sex differences within a strain.

Similar to the analysis of acquisition of oxycodone intake, the mean intake during the first three escalation sessions (“early”) and last three escalation sessions (“late”) was calculated **(Supplemental Figure 3)**. Oxycodone intake was influenced by a strain-sex (F_13,184_ = 3.83, p < 0.001) and a strain-timepoint (F_13,184_ = 3.46, p < 0.001) interaction suggesting that sex differences were not consistent across strains and the magnitude of escalation was not consistent across strains. To further describe strain differences the change in oxycodone intake during the escalation phase was calculated **(Figure 7)**. A main effect of strain (F_13,184_ = 3.48, p < 0.001) highlights the significant variation between the strains in their propensity to increase oxycodone intake during the escalation phase.

**Figure 7.**
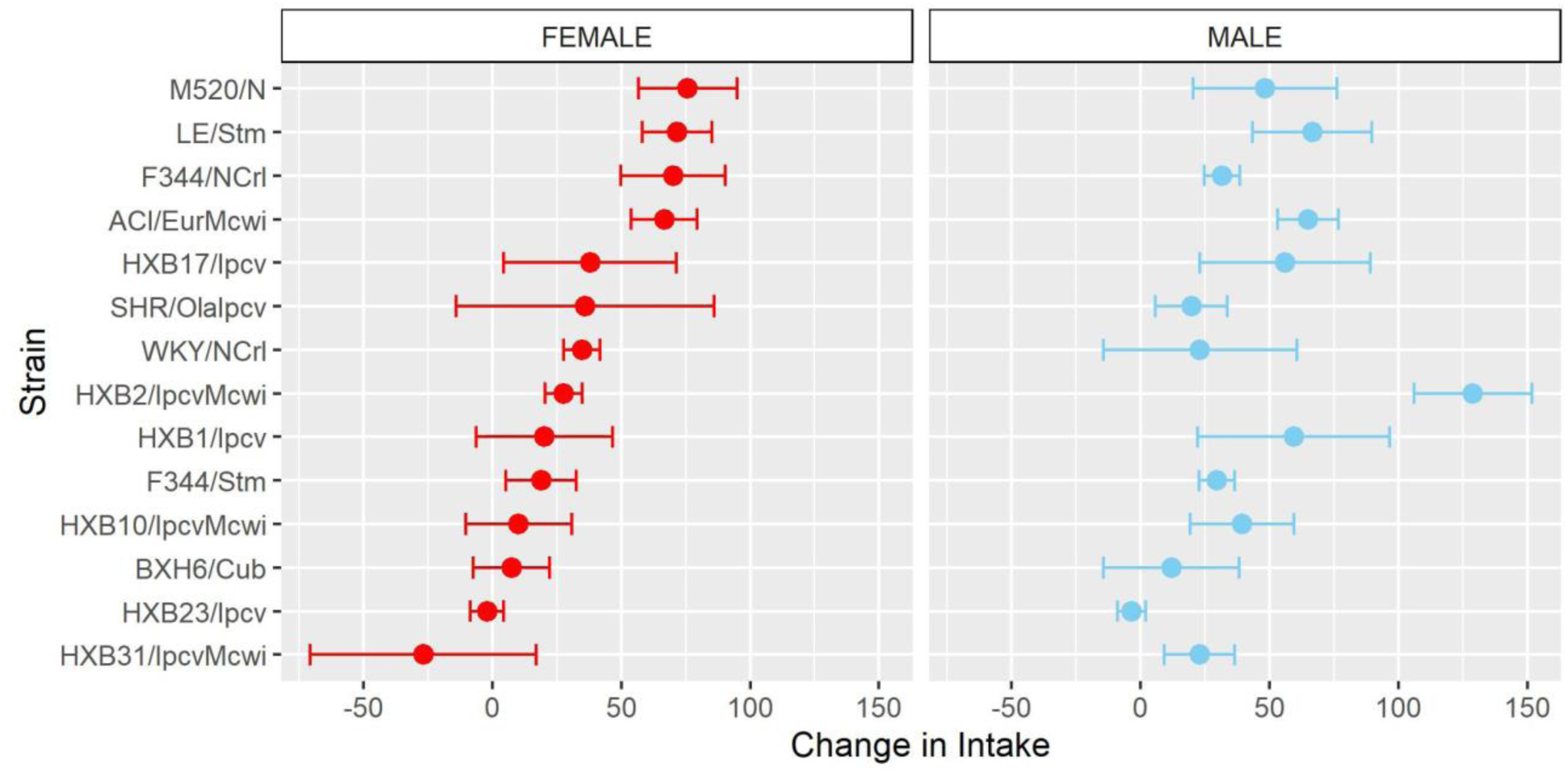
Change in oxycodone intake during escalation is influenced by genetic background. The change in oxycodone intake was determined by calculating the difference between early and late escalation sessions. The mean ± SEM for each strain is plotted for females (left) and males (right). Strains for both females and males are displayed in order of increasing change in females. A main effect of strain influenced the change in oxycodone intake across the escalation phase.

### 3.3 Lever discrimination

As another measure of behavioral responding during the oxycodone self-administration procedure, we calculated each rat’s ability to discriminate between the lever paired with oxycodone delivery and the inactive lever. Lever discrimination was evaluated by calculating the ratio of correct responses to total responses for each rat within individual sessions. Therefore, a value close to one indicates that the rat is almost exclusively pressing the active lever. To evaluate how lever discrimination changes across sessions, we calculated the mean lever discrimination during the first three sessions (“early”) and last three sessions (“late”) of the acquisition and escalation phases. During acquisition **(Figure 8A)**, lever discrimination was shaped by strain-sex (F_14,236_ = 2.31, p = 0.005) and strain-timepoint (F_14,236_ = 3.44, p < 0.001) interactions. During escalation **(Figure 8B)**, lever discrimination was influenced by a strain-timepoint interaction (F_13,184_ = 2.08, p = 0.017). These findings suggest that learning to discriminate the outcomes between the two levers generally improves or stays the same throughout the procedure but is also dependent on the strain and sex of the animal. It is interesting that rats from the genetically related SHR/OlaIpcv and WKY/NCrl strains (36) displayed divergent patterns of lever discrimination across both phases of self-administration **(Figure 8C)**. Lever discrimination in WKY/NCrl rats improved throughout both phases of self-administration while SHR/OlaIpcv rats show persistent deficits in lever discrimination hovering around 50% lever discrimination.

**Figure 8.**
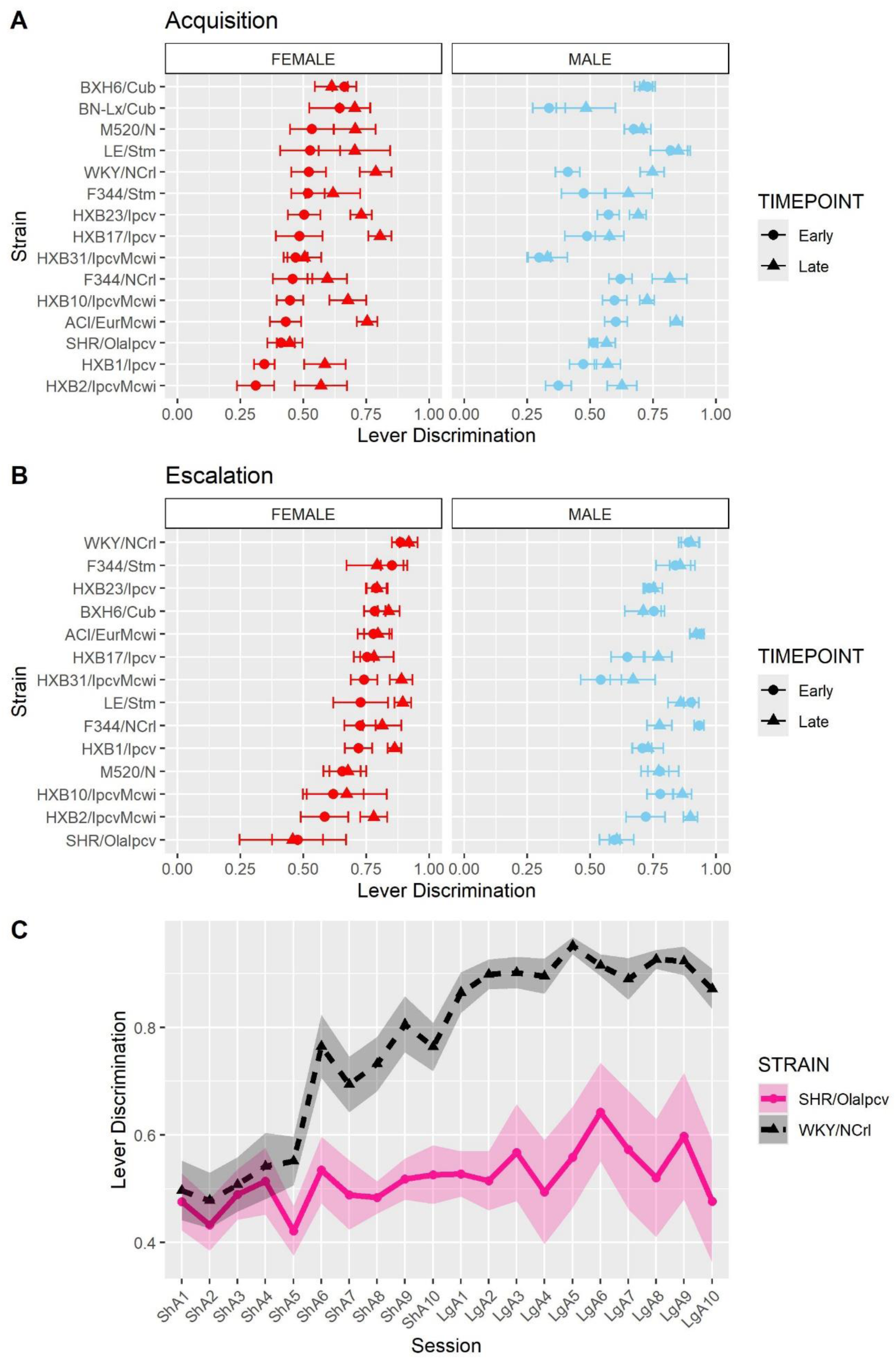
Lever discrimination is influenced by strain and sex. Lever discrimination was calculated as the ratio of correct lever responses to total responses within each session. **(A)** The mean ± SEM lever discrimination for each strain during acquisition is plotted for females (left) and males (right). Strains for both females and males are displayed in order of increasing discrimination during the early timepoint in females. Lever discrimination during acquisition was shaped by a strain-sex and a strain-timepoint interaction. **(B)** The mean ± SEM lever discrimination for each strain during escalation is plotted for females (left) and males (right). Strains for both females and males are displayed in order of increasing discrimination during the early timepoint in females. During escalation, lever discrimination was influenced by a strain-timepoint interaction. **(C)** Strikingly divergent patterns of lever discrimination were observed in rats from two genetically related strains (SHR/Olalpcv and WKY/NCrl strains).

### 3.4 Normalized timeout responding

We also evaluated the persistence of responding during periods of drug unavailability by evaluating drug-paired lever responding during the 20-s timeout period following each oxycodone infusion. Because animals that self-administer more oxycodone have more opportunities for timeout responding, the number of timeout responses was normalized to the number of oxycodone infusions. Normalized timeout responding during acquisition **(Figure 9A)** was influenced by a strain-sex-timepoint interaction (F_14,236_ = 1.78, p = 0.043). Although most animals displayed low levels of timeout responding throughout acquisition, it was striking that some strains (e.g., HXB1/Ipcv) were more likely to engage in persistent responding during the timeout period, especially during the late timepoint. Normalized timeout responding during escalation **(Figure 9B)** was influenced by a strain-sex (F_13,184_ = 3.79, p < 0.001) and a strain-timepoint (F_13,184_ = 3.11, p < 0.001) interaction. Here, males from a few strains (e.g., HXB2/IpcvMcwi) displayed a robust increase in timeout responding.

**Figure 9.**
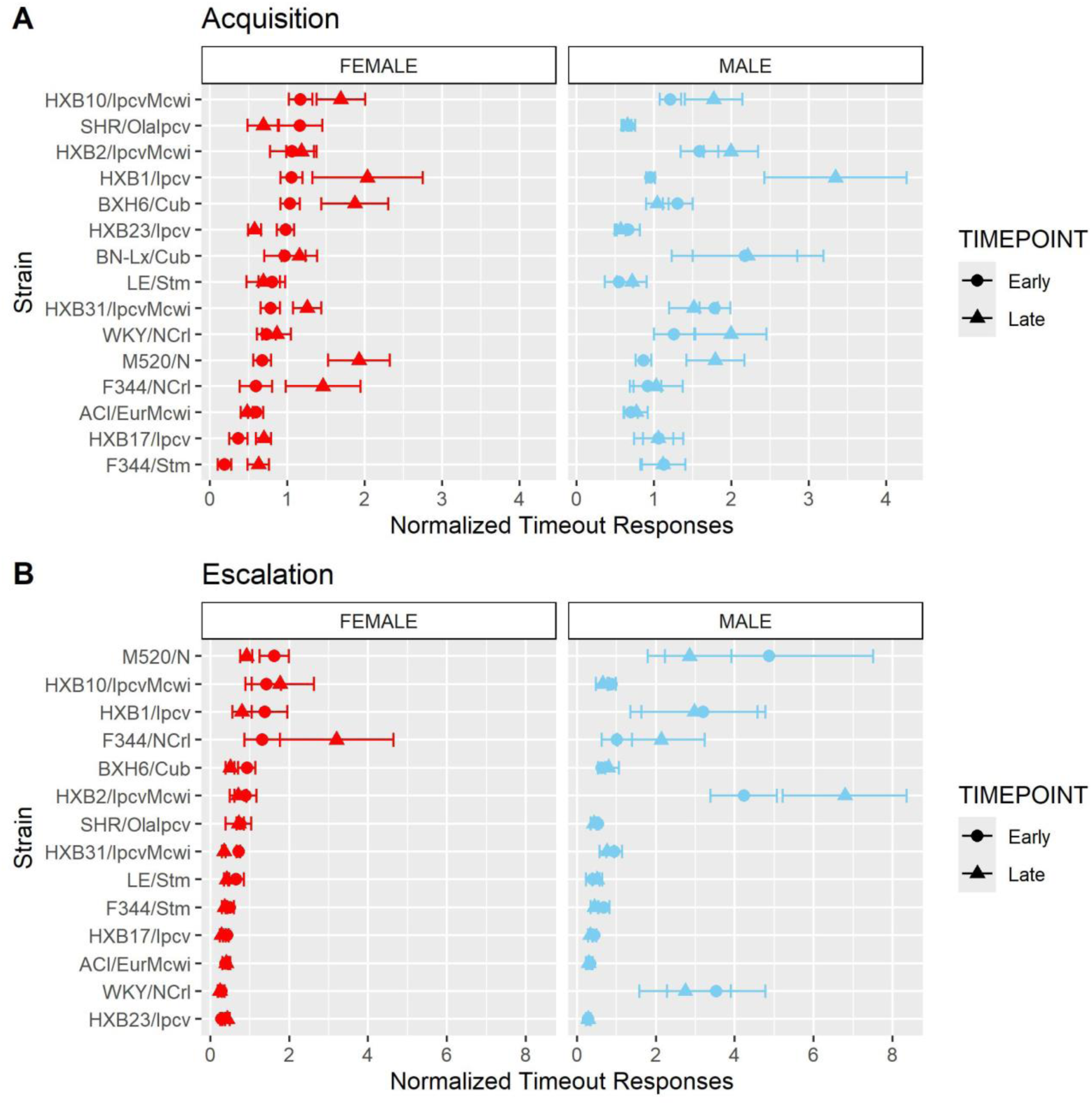
Normalized timeout responding is influenced by strain and sex. The number of timeout responses was normalized to the number of infusions within sessions. **(A)** The mean ± SEM normalized timeout responding for each strain during acquisition is plotted for females (left) and males (right). Strains for both females and males are displayed in order of increasing timeout responding during the early timepoint in females. During acquisition, normalized timeout responding was influenced by a strain-sex-timepoint interaction. **(B)**. The mean ± SEM normalized timeout responding for each strain during escalation is plotted for females (left) and males (right). Strains for both females and males are displayed in order of increasing timeout responding during the early timepoint in females. During escalation, normalized timeout responding was influenced by a strain-sex and a strain-timepoint interaction.

### 3.6 Correlations between oxycodone-related phenotypes

To evaluate the relationships between oxycodone-related behavioral phenotypes, Pearson correlations were calculated using strain means within each sex (**Figure 10)**. Phenotypes related to oxycodone-induced analgesia, as described previously in (31), were included to describe the relationship between oxycodone analgesia and self-administration phenotypes. The maximal possible efficacy (% MPE) was used as a measure of acute analgesia 15- and 30-minutes after oxycodone administration, where higher values represent a greater degree of oxycodone-induced analgesia **(Supplemental Methods**). All pairwise comparisons between phenotypes are included in **Supplemental Table 3**. Select correlations between phenotypes are displayed in **Figure 11**. The magnitude of oxycodone-induced analgesia at 30-minutes positively correlated with total oxycodone intake during escalation (r(26) = 0.39, p = 0.04) **(Figure 11A)**. This suggests that drug-naïve animals that experience a greater degree of acute oxycodone-induced analgesia may go on to consume higher levels of oxycodone. The normalized timeout responding during early acquisition sessions negatively correlated with lever discrimination during late acquisition sessions (r(26) = −0.42, p = 0.02) **(Figure 11B)**. This suggests that high levels of responding during a period of drug unavailability correlate with poor lever discrimination during the acquisition phase.

**Figure 10.**
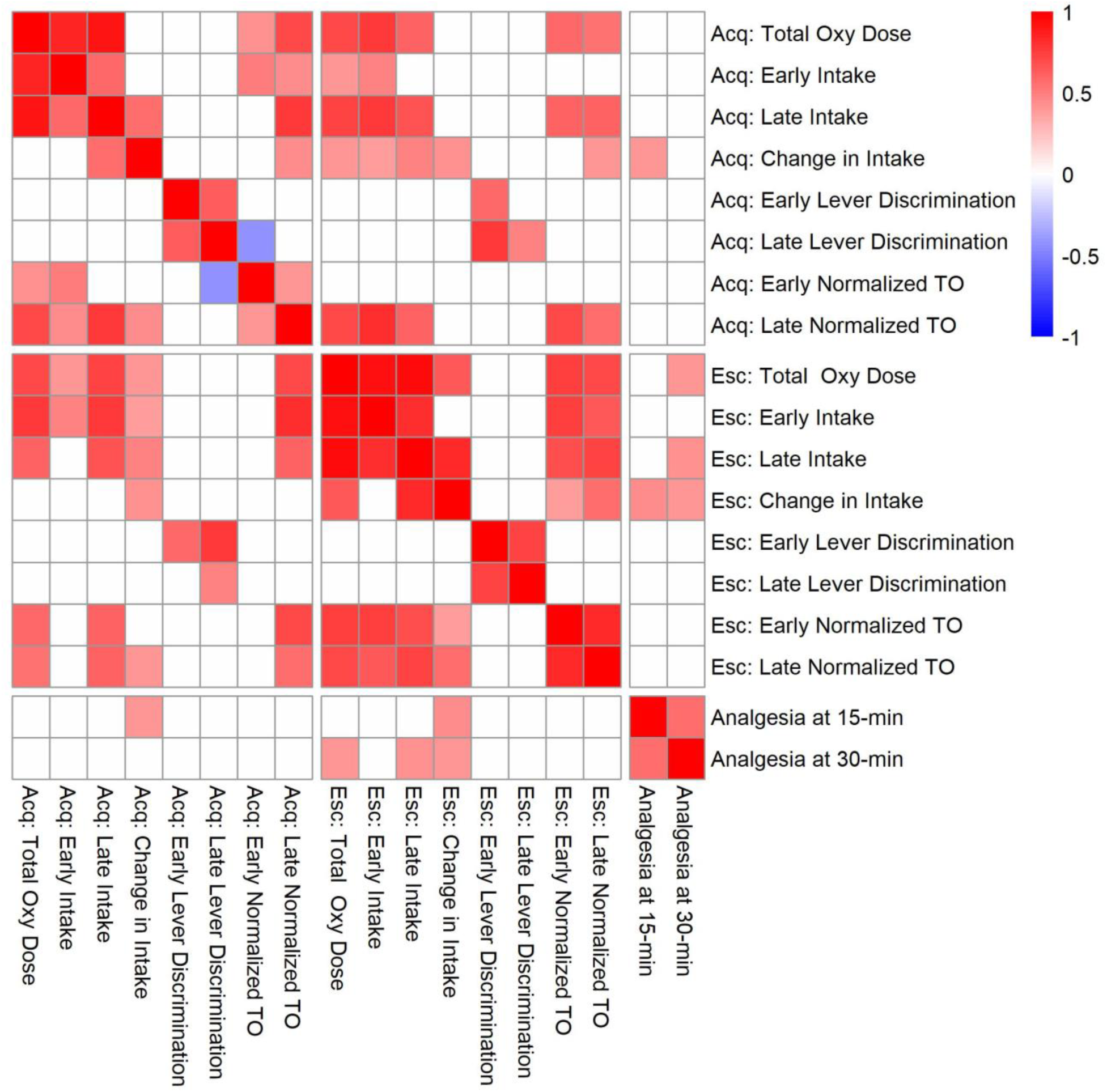
Correlation matrix for analgesia phenotypes and oxycodone self-administration phenotypes during the acquisition (Acq) and escalation (Esc) phases. Strain means calculated within each sex were used to evaluate genetic correlations between oxycodone self-administration and analgesia phenotypes. Only significant correlations (p < 0.05) are shaded, and white boxes indicate no significant correlation. See Supplemental Table 3 for p-values and Pearson’s r values for all comparisons.

**Figure 11.**
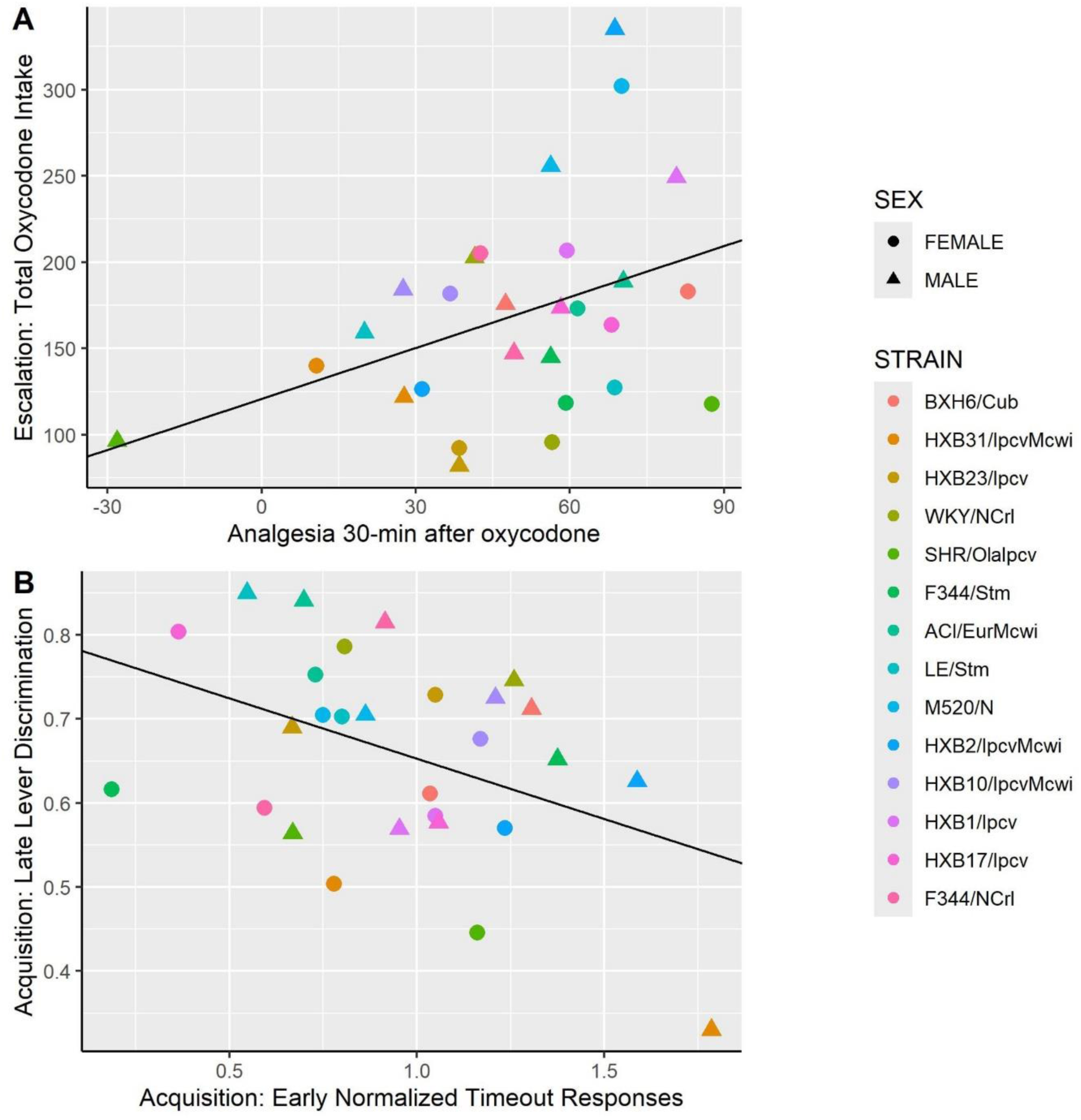
Correlations between select oxycodone phenotypes. **(A)** The degree of oxycodone-induced analgesia (% MPE) 30-minutes after an oxycodone injection (1 mg/kg, i.p.) was positively correlated with the total dose of oxycodone self-administered during escalation. **(B)** During acquisition, the number of normalized timeout responses in early sessions was negatively correlated with lever discrimination in late sessions.

### 3.7 Heritability estimates

The broad-sense heritability (H^2^) of several behavioral phenotypes was calculated by a 2-way ANOVA using strain and sex as the independent variables **(Table 2)**. Measures of oxycodone intake were highly heritable, ranging between H^2^ = 0.26-0.54. Other aspects of oxycodone self-administration, such as lever discrimination and normalized timeout responding, were more moderately heritable, ranging between H^2^ = 0.25-0.42.

**Table 2:** Broad-sense heritability estimates for self-administration phenotypes during the acquisition and escalation phases. Heritability estimates were calculated using the coefficient of determination (R^2^) from a 2-way ANOVA with strain and sex as the predictors.

| Phenotype | Acquisition Heritability | Escalation Heritability |
| --- | --- | --- |
| Total Oxycodone Intake | 0.50 | 0.54 |
| Early Oxycodone Intake | 0.48 | 0.33 |
| Late Oxycodone Intake | 0.42 | 0.48 |
| Change in Intake | 0.32 | 0.26 |
| Early Lever Discrimination | 0.29 | 0.31 |
| Late Lever Discrimination | 0.27 | 0.25 |
| Early Normalized Timeout Responses | 0.34 | 0.35 |
| Late Normalized Timeout Responses | 0.33 | 0.42 |

## Discussion

Individual genetic variation plays a major role in the risk of using and misusing opioids (12,37). Using selected inbred rat strains from the HRDP, we demonstrated that sex and genetic background influence several specific features of oxycodone self-administration. We observed that all male and female rats reliably acquire oxycodone self-administration, and genetic background and sex contribute to variability in the escalation of oxycodone intake. Other measures observed during oxycodone self-administration, such as lever discrimination and timeout responding, were also mediated by interactions between strain and sex. Nearly all measures of oxycodone intake were highly heritable (H^2^ = 0.26-0.54), while the measures of learning (lever discrimination) and drug seeking (timeout responding) were slightly less heritable (H^2^ = 0.25-0.42).

We found that the interaction between genetic background and sex contributes to behavioral variance in oxycodone self-administration. This complements previous findings that oral oxycodone self-administration is highly heritable and that sex differences in self-administration are dependent on genetic background (32,33). Sex differences have not consistently been observed in previous oxycodone self-administration studies (32,38–42). Several inbred and outbred rat strains do not exhibit sex differences in the pattern of i.v. oxycodone self-administration (38,39,42). In the limited number of strains that do exhibit sex differences, the directionality of the sex difference is inconsistent, potentially due to differences in fixed-ratio schedules and the oxycodone dose per infusion (32,40,41). Together, this indicates that sex differences in oxycodone self-administration are highly dependent on genetic background and the specific self-administration paradigm. We observed that WKY/NCrl males display nominally higher levels of oxycodone intake than females. Interestingly, the interaction between sex and genetic background influences oxycodone pharmacokinetics (32). WKY male rats display higher peak levels of plasma oxycodone after an intravenous infusion compared to WKY females, which may contribute to the nominal sex differences in oxycodone intake we observed.

Our lab previously demonstrated that sex and genetic background contribute to the magnitude and duration of oxycodone-induced analgesia (31). In the current study, the efficacy of oxycodone-induced analgesia in drug-naïve animals was positively correlated with oxycodone intake during the escalation phase of self-administration. This suggests that animals having greater sensitivity to oxycodone-induced analgesia also consume higher amounts of oxycodone. Previous work in male Sprague-Dawley rats indicates that low sensitivity to morphine-induced analgesia is associated with the escalation of morphine intake (43). This discrepancy may be due to the pharmacokinetic and pharmacodynamic properties of the opioids being tested or genetic differences between strains. For example, it is well known that stimulation of the mu-receptor mediates analgesia and reward-related phenotypes (44,45). Genetic variants in the *Oprm1* gene that impact mu receptor function may underlie the relationship between these behavioral phenotypes. A single nucleotide polymorphism in the *Oprm1* gene in mice alters both opioid-induced analgesia and self-administration (46–49). Strain differences in mu-receptor polymorphisms, gene or protein expression, or alterations in protein function may modulate this variability in analgesia and self-administration phenotypes.

Genetic background contributed to variability in lever discrimination throughout the oxycodone self-administration procedure. Two genetically related strains, the Spontaneously Hypertensive Rats (SHR) and Wistar-Kyoto rats (WKY), displayed divergent patterns of lever discrimination throughout the self-administration period. The SHR strain has traditionally been used as a rodent model of attention deficit hyperactivity disorder and is often compared with the WKY strain when assessing learning and memory phenotypes (50–53). In our experiment, WKY/NCrl rats increased lever discrimination throughout the acquisition phase and displayed stable, high levels of lever discrimination during the escalation phase. SHR/OlaIpcv rats, however, showed low levels of lever discrimination that did not improve over time. This may suggest that SHR rats display deficits in learning to distinguish the outcomes resulting from lever responding. Poor lever discrimination may also be influenced by other related and/or competing traits. Male SHR rats also display greater levels of locomotion than WKY rats in the locomotor, open field, and elevated plus maze tasks (54,55). Thus, the poor lever discrimination in SHR rats may result from generalized psychomotor activation that causes indiscriminative responding at both levers. Oxycodone-induced locomotor activity and sensitization are heritable phenotypes, and these strains display may differ in these traits further contributing to poor lever discrimination (56). Finally, SHR rats also display greater impulsivity in non-drug learning procedures, suggesting that higher impulsivity could contribute to deficits in lever discrimination (50,53,57,58).

We observed that interactions between strain and sex influenced normalized timeout responding during the acquisition and escalation phases. During the escalation phase, males from several strains displayed higher levels of timeout responding than female counterparts. Interestingly, outbred Sprague-Dawley males also display high levels of timeout responding during oxycodone self-administration (38). Drug-seeking that continues during periods of drug unavailability has been associated with compulsive drug use patterns and may suggest that some strains are more prone to this behavioral phenotype (59). Responding during the timeout period may also indicate perseverative responding with higher levels of timeout responding associated with an increased effort to try to obtain oxycodone (60,61). Timeout responding may also represent a learning deficit, where animals fail to associate the activation of the cue light with the period of drug unavailability. It will be important to conduct more precise measurements to tease out these interpretations and more accurately interpret these findings.

Together, these results suggest that genetic variance between strains contributes to variability in several aspects of oxycodone self-administration. This population of rats is an ideal candidate for future genetic mapping studies and can be effectively used to examine how the body and brain respond to long-term oxycodone use. We plan to examine how strain-based variation in oxycodone intake alters the gut microbiome and gene expression in the brain.

## Data Availability

Data are available on request from the authors.

## Acknowledgements

The authors thank Melinda Dwinell PhD (Medical College of Wisconsin; R24OD024617) for providing several of the inbred strains studied in this article. We thank Caleb Hodges and Safa Vaseemuddin for assistance with collecting behavioral data.

