## Supplemental Methods & Figures for "Sex and genetic background influence intravenous oxycodone self-administration in the Hybrid Rat Diversity Panel"

#### Tail Immersion Test for Thermal Sensitivity and Oxycodone-Induced Analgesia

Prior to surgical and self-administration procedures, animals underwent testing in the von Frey test of mechanical sensitivity and the tail immersion test of thermal sensitivity to evaluate somatosensation and oxycodone-induced analgesia (1). Briefly, 7-8 days prior to surgery, animals underwent the tail immersion test to examine thermal sensitivity and oxycodone-induced analgesia.

The tail immersion test was used to measure thermal sensitivity and the analgesic response to oxycodone. During each trial, the distal 5-6 cm of the rat's tail was immersed in 49 ± 0.5 °C water. The latency for a tail flick or tail withdrawal response was recorded. A cutoff of 15 seconds (s) was used to prevent tissue damage. These parameters were established based on preliminary studies identifying that some strains experienced greater tissue damage at prolonged immersion times (>15 s) and/or higher temperature intensities (>50°C). Baseline tail withdrawal latencies were recorded prior to oxycodone administration (1 mg/kg, i.p.) to determine the strain differences in thermal sensitivity.

Oxycodone-induced analgesia was measured by recording tail withdrawal latencies 15-, 30-, 45-, 60-, 90-, and 120-minutes following oxycodone administration (1 mg/kg, i.p.). The maximum possible efficacy (% MPE) was used as a measure of acute analgesia 15- and 30-minutes after oxycodone administration. % MPE was calculated as follows:

$$[(\text{Latency}_x - \text{Baseline Latency}) / (\text{Cutoff Latency} - \text{Baseline Latency})] \times 100\%.$$

#### Outlier Detection and Winsorization

Prior to statistical analysis, outliers were detected and winsorized. For self-administration behaviors, the responses (correct, incorrect, and timeout) were summarized for both sexes within each strain for each session during the acquisition and escalation phases. Outliers were defined as daily responses (summarized using strain, sex, and session) greater than Q3 + 3IQR or less than Q1 - 3IQR, then winsorized to the outlier detection threshold. A similar procedure was performed for oxycodone-induced analgesia (% MPE) 15- and 30-minutes after an oxycodone injection.

#### Missing Data Imputation

Missing session data (< 2% of self-administration session data) was imputed within each rat. If the missing session occurred during the middle of a self-administration phase, the data was imputed by calculating the mean from the session before and the session after the missing session. If the missing session occurred on the last day of the self-administration phase, the data was imputed by calculating the mean from the previous two sessions before the missing session.

### Supplemental Figures

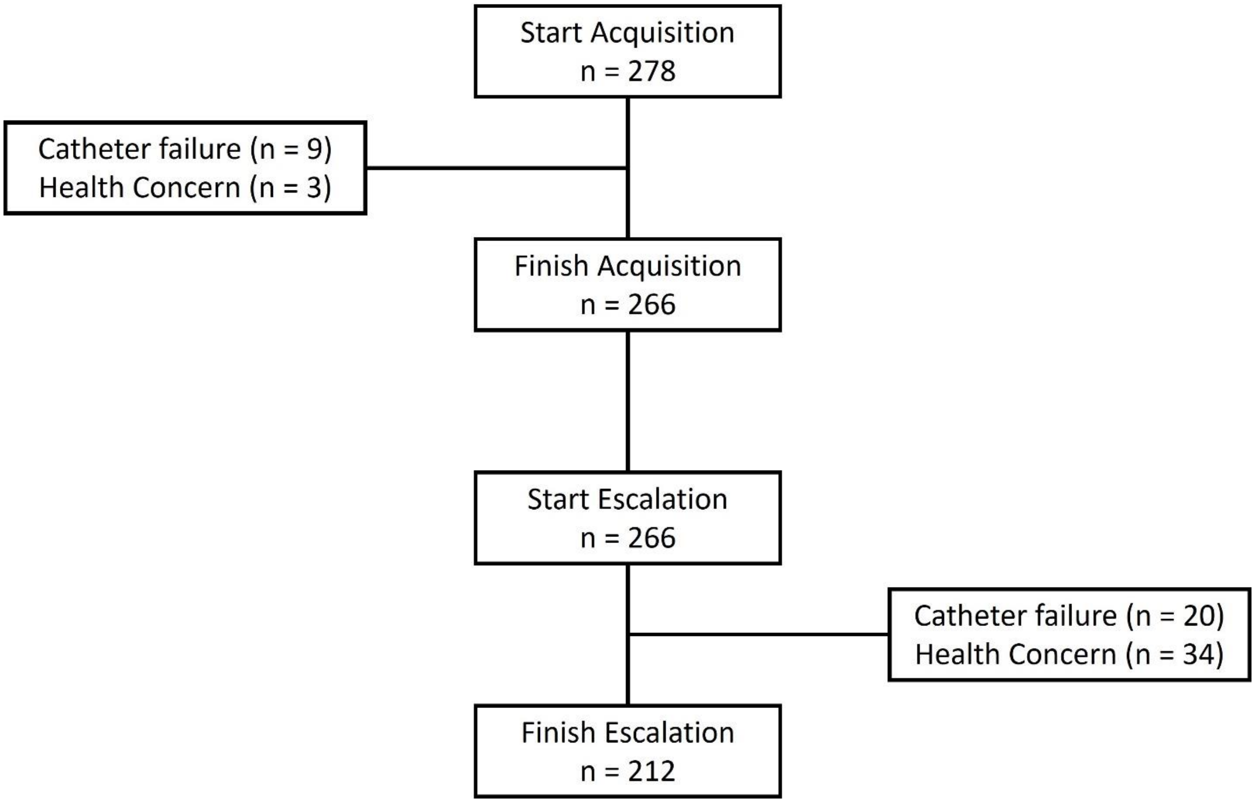


**Supplemental Figure 1.** Flow chart displaying total sample sizes and reasons for attrition throughout the acquisition and escalation phases. Animals were excluded from the study if their catheter failed following a propofol test or for potential health concerns (e.g., weight loss, overdose).

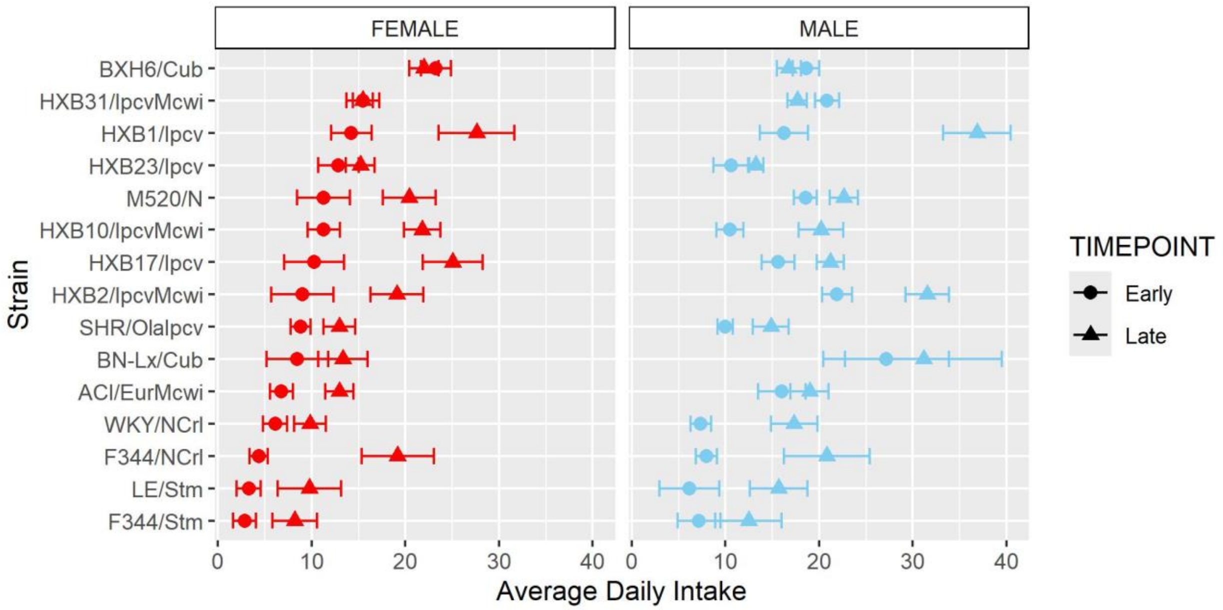


**Supplemental Figure 2.** Oxycodone intake during early and late acquisition sessions. During acquisition, intake was shaped by strain-sex and strain-timepoint interactions. Strains for both females and males are displayed in order of increasing average daily intake during the early timepoint in females.

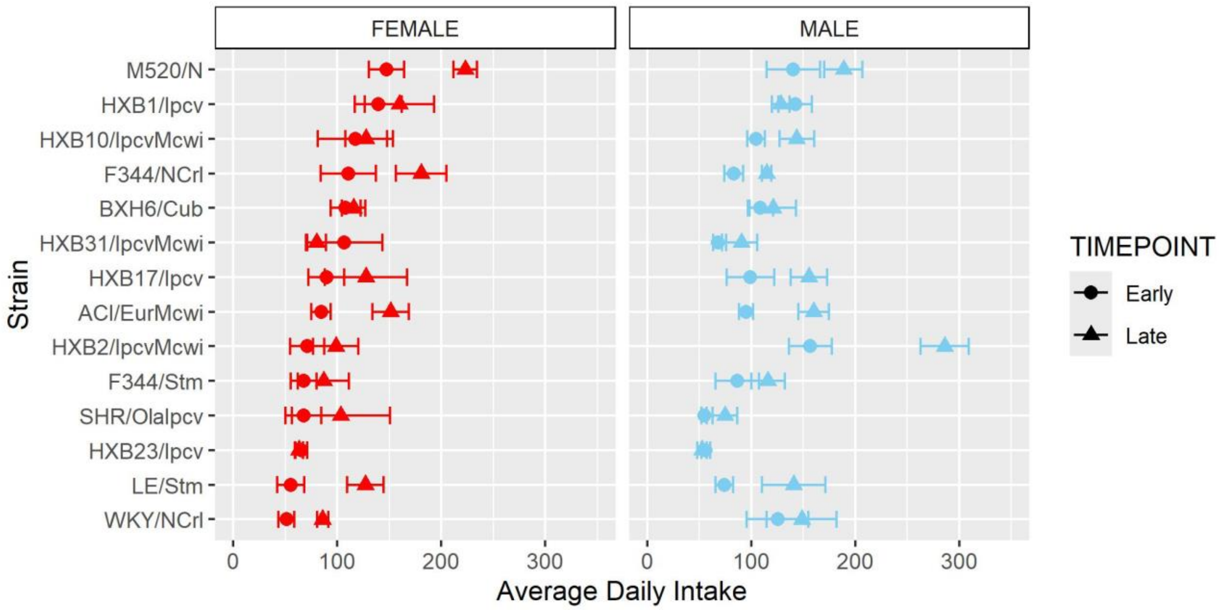


**Supplemental Figure 3.** Oxycodone intake during early and late escalation sessions. During escalation, oxycodone intake was influenced by strain-sex and strain-timepoint interactions. Strains for both females and males are displayed in order of increasing average daily intake during the early timepoint in females.
